# Identification of critical factors of the replication stress response in human cells

**DOI:** 10.1101/2025.01.12.632149

**Authors:** Louise M.E. Janssen, Empar Baltasar Perez, Chantal Vaarting, Abdelghani Mazouzi, Matthijs Raaben, Thijn R. Brummelkamp, René H. Medema, Jonne A. Raaijmakers

**Affiliations:** Division of Cell Biology, Oncode Institute, The Netherlands Cancer Institute, Plesmanlaan 121, 1066 CX Amsterdam, The Netherlands; Division of Biochemistry, Oncode Institute, The Netherlands Cancer Institute, Plesmanlaan 121, 1066 CX Amsterdam, The Netherlands; Oncode Institute, Princess Maxima Center for Pediatric Oncology, Heidelberglaan 25, 3584 CS Utrecht, The Netherlands

## Abstract

High fidelity of replication is important to preserve genomic integrity and ensure healthy progeny. Perturbations of replication, also known as replication stress, is frequently observed in cancer cells and is considered a cancer cell-specific trait. Although replication stress drives genomic instability and tumor progression, it also generates a targetable cancer-specific vulnerability. In order to identify potential therapeutic targets in cancer cells that experience replication stress, we performed a genome wide genetic screen in human HAP1 cells challenged with low doses of replication stress-inducing drugs. We identified a large set of genes that specifically hamper cell survival in the context of replication stress. In addition to well-known players in the replication stress response and DNA repair, such as RNASEH1, BRIP1, and MDC1, we identified several genes with no prior described role in DNA replication, damage tolerance or repair. We validated that the loss of GIGYF2, HNRNPA2B1, and SUMO2 renders cells more vulnerable to replication stress. For GIGYF2 and SUMO2, we could implicate a role in homologous recombination. Taken together, our replication stress screen identified several known as well as some novel factors that protect against the toxic implications of replication stress. These factors could entail potential therapeutic targets for cancer cells experiencing replication stress.

## Introduction

Proper DNA replication is a vital process for cell division, as faithful duplication of the genome is a key step to ensure cell survival and to prevent malignant transformation^1–3^. Replication is an orchestrated process but can be challenged by both endogenous and exogenous causes, such as a limited deoxynucleoside triphosphates (dNTP) supply, collisions with transcription, and DNA damage. Such insults can cause the replication fork to slow down or stall completely^1^. These impediments of replication are altogether referred to as replication stress (RS)^1^. Fork delay or stalling creates stretches of single-stranded DNA (ssDNA), which are recognized by replication protein A (RPA)^1,4^. Upon binding to ssDNA, RPA promotes the activation of the RS-response via activation of ATR, which results in delayed S-phase progression, thereby providing time to resolve the issues and limit error-prone genome duplication^1,5,6^.

Under normal conditions, replicating cells display low levels of RS^7,8^. However, the activation of oncogenes in precancerous lesions can induce persistent RS, which results in increased genomic instability^1,9–11^. Oncogene activation drives replications stress via a plethora of mechanisms^12^. For example, amplification of the oncogenes *MYC* or *CCNE1* causes a loss of cell cycle control, resulting in a shortened G1 phase and a failure to license sufficient replication origins^11^. Additionally, it has been shown that overexpression of oncogenes, such as KRAS or E7, results in reduced fork speed due to lower dNTP availability. Oncogene activation often results in a failure to coordinate transcription and replication, thereby inducing collisions that lead to fork collapse and DNA breaks at fragile sites. Finally, oncogenes or tumor suppressors that affect the DNA damage response, cell cycle checkpoints or DNA damage repair can also result in RS in a more indirect manner. The genomic instability caused by oncogene-induced replication stress results in the accumulation of additional mutations and chromosomal aberrations that drive tumor heterogeneity and cancer progression ^13–15^.

The fact that RS is a cancer cell-specific hallmark creates an opportunity to identify cancer-specific vulnerabilities^7,16,17^. Here, we aimed to identify such vulnerabilities using a genetic screen in cells treated with RS-inducing drugs. Using this unbiased approach, we identified several genes with a known role in the RS response but also a number of genes with no previously established role in the RS-response or in DNA damage repair to be specifically required in cells suffering from RS. This list of genes contains a large group of mitochondrial-related genes, a translation regulator *GIGYF2*, an RNA binding protein HNRNPA2B1 and the ubiquitin-like gene *SUMO2*.

## Results

### Induction of replication stress in haploid human cells

In order to identify vulnerabilities of cells experiencing RS, we performed genetic synthetic lethality screens in human haploid cells. In these screens, we induced RS by partially inhibiting enzymes involved in replication; either the DNA polymerase-α using Aphidicolin (Aph)^18^, or the ribonucleotide reductase using hydroxyurea (HU)^19^, thereby reducing the cellular dNTP pool. Both of these methods were previously shown to inhibit the replication machinery and result in delayed replication fork progression^20^. To test the effect of these drugs on HAP1 cell viability, we performed growth assays in which we treated wild-type (WT) HAP1 cells with increasing doses of either Aph or HU (Fig. 1A) and calculated an IC50 of 66 nM for Aph, and an IC50 of 140 μM for HU (Fig. 1B).

**Figure 1:**
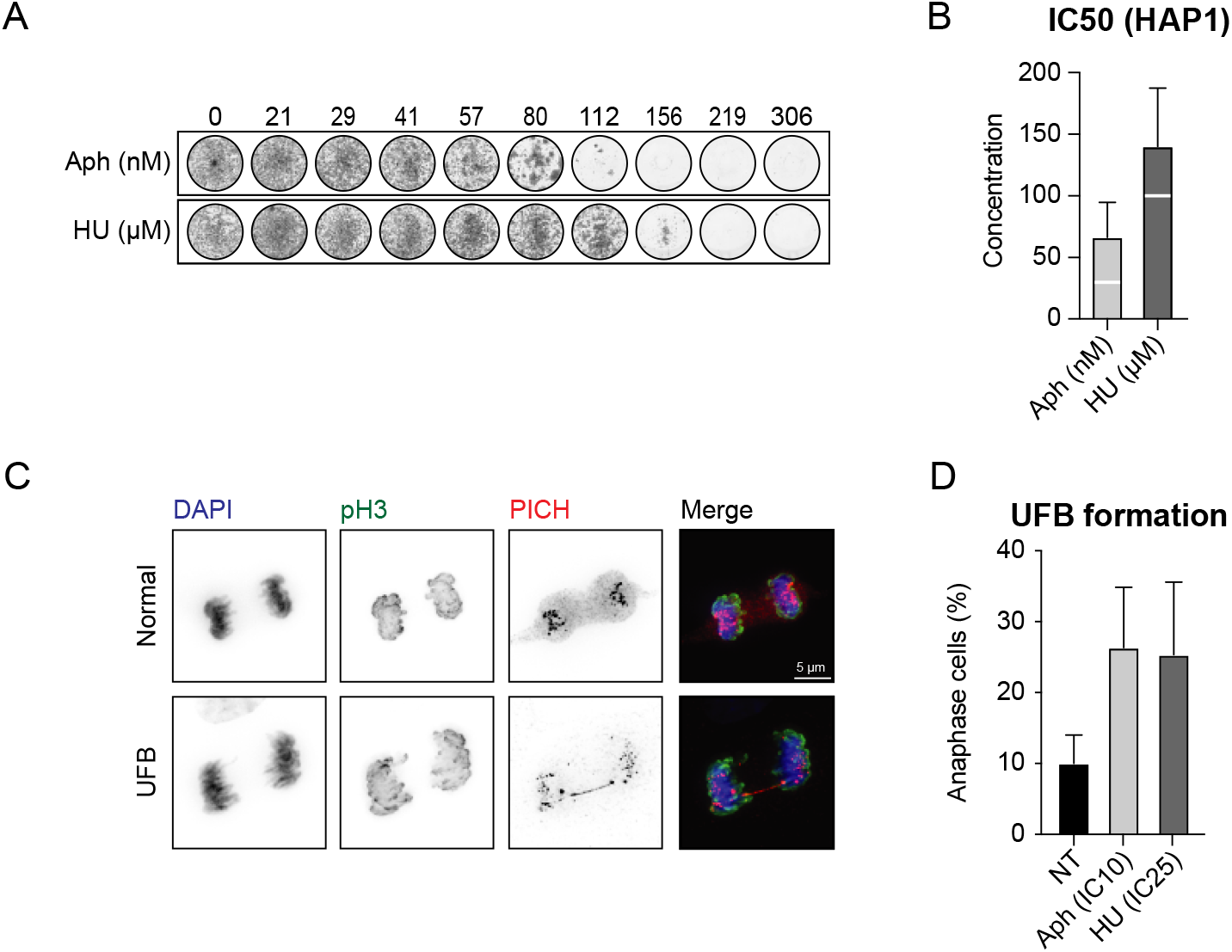
Induction of replication stress in HAP1 cells. **(A)** Representative growth assay of HAP1 cells subjected to increasing doses of Aph or HU for 5 days. **(B)** Average IC50 of 5 independent growth assays (IC50 for Aph 66 nM, for HU 140 μM). White line represents IC10 (Aph) or IC25 (HU). **(C)** Representative images of immunofluorescent staining of PICH fibers in anaphase HAP1 cells. **(D)** Quantification of PICH fibers in anaphase HAP1 cells treated with 27.5 nM Aph (IC10) or 100 μM HU (IC25) for 48 hours. Average of 4 independent experiments, mean + SD, differences just below significance.

Despite the fact that cells have mechanisms in place to detect RS and induce a checkpoint response^21–26^, mild RS is shown to be tolerated to a certain extent^27^. This causes cells to progress into mitosis with under-replicated DNA. This can create unstable DNA structures like ultra-fine bridges (UFBs), ultimately leading to DNA damage^13,28–31^. Because we aimed to induce chronic RS for multiple days, we set out to identify a drug concentration where cells show signs of RS-induced genomic instability, while proliferation was minimally affected. For this, we measured the occurrence of PICH fibers, a readout of UFBs^30,32^, by immunofluorescence (Fig. 1C). Untreated WT HAP1 cells showed a baseline level of 10% anaphase cells with UFB’s (Fig. 1D), comparable to previous reports^33,34^. When treating HAP1 cells with low doses of Aph or HU (using the IC10 of 27.5 nM, or the IC25 of 100 μM respectively) we were able to elevate the percentage of the anaphase cells with UFBs to ∼25% (Fig. 1D), yet cells were still able to proliferate over multiple days (Fig. 1A).

### A synthetic lethality screen identified genes essential for survival under replication stress

Using Aph and HU to induce RS-dependent genomic instability, we set out to discover what genes are important for cell viability in cells that experience RS. To this end, we performed genome-wide insertional mutagenesis in near-haploid HAP1 cells and subsequently treated the cells with for 10 days with the low doses of HU and Aph established in Figure 1 (see ^35^ and **methods** for more details). We performed a single replicate for cells treated with a low dose HU and two independent replicates of cells treated with a low dose of Aphidicolin. Genes were considered hits when they significantly dropped out in HU and in at least one of the Aphidicolin replicates to avoid drug-specific interactions. The hits displayed a significant correlation between all different screens, reflecting a high reproducibility (Fig S1). In total, we identified 81 genes to be synthetic lethal with RS (Fig. 2A-D).

**Figure 2:**
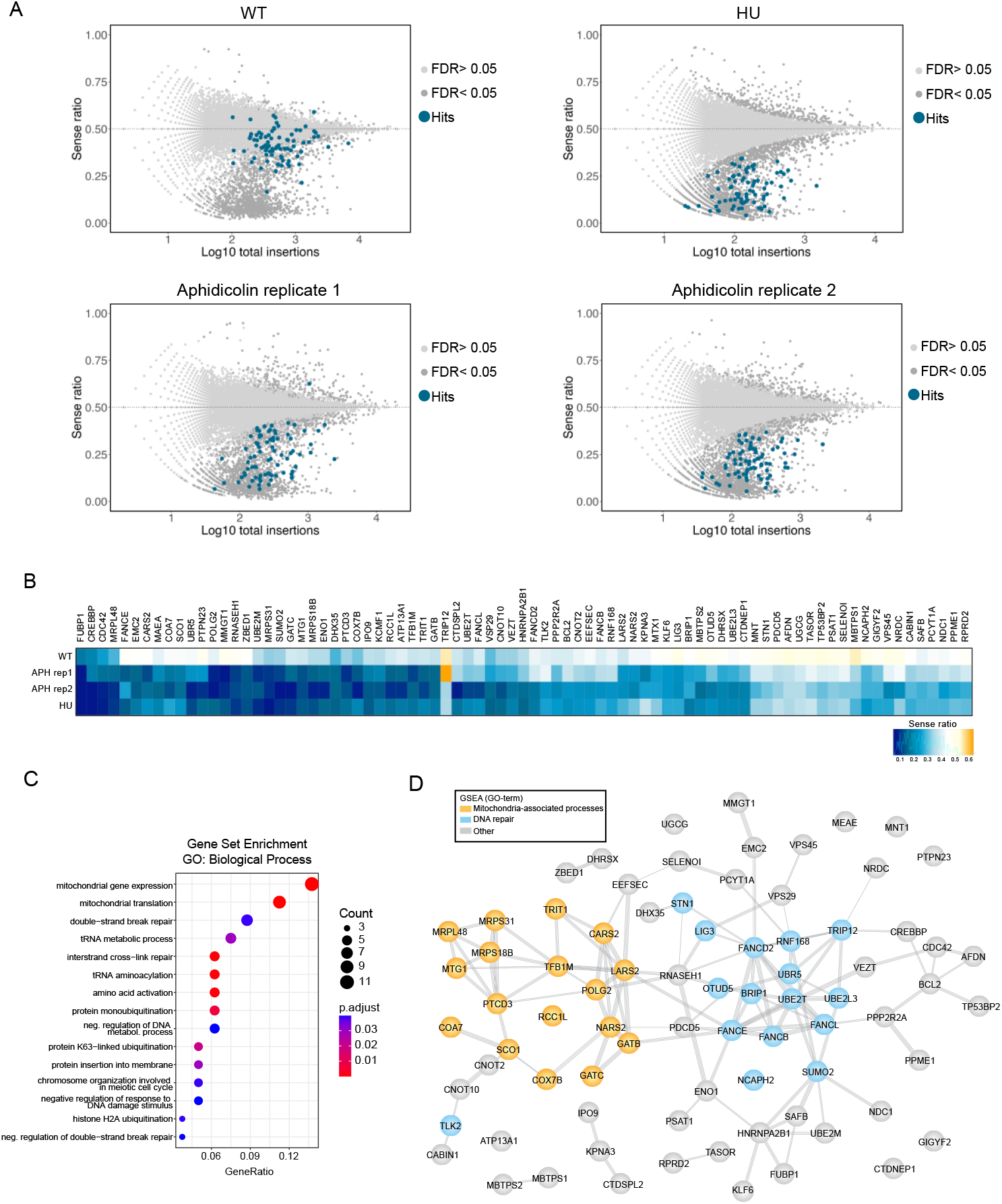
Synthetic lethality screen in HAP1 cells identifies factors essential for survival during replication stress. **(A)** Fishtail plots of gene-trap insertions in WT HAP1 cells (representative example), 2 replicates of HAP1 cells treated with a low dose of Aphidicolin and 1 replicate of cells treated with a low dose of HU. Each dot in the plot represents a single gene, and the number of integrations per gene is plotted on the x-axis. On the y-axis, the relative amount of sense integrations over the total amount of integrations (sense+antisense) is plotted. Genes were considered hits if they were significantly selective in each individual replication stress condition (Binomial FDR <0.05), significantly different from 4 individual WT samples (p<0.05) and a subtraction ratio (ratio WT – ratio replication stress) >0.12. Moreover, the genes needed to overlap in the HU condition and at least one of the Aphidicolin replicates. There were 81 genes passing these criteria (hits, highlighted in blue) **(B)** Heatmap displaying the average ratio of sense/total integrations in WT (average of 4 screens), HU or APH treated HAP1 cells. Graph contains the hitlist consisting of 81 genes. **(C)** Gene set enrichment analysis for GO terms “Biological Process” for the 81 hits **(D)** STRING interaction network of the hits (B). Hits are color coded based on their previously described roles.

The RS response and DNA damage response (DDR) are closely intertwined. Stalled forks exhibit large stretches of single stranded DNA which result in the recruitment of DDR proteins such as RPA and ATR that trigger a DNA damage response and result in the recruitment of several downstream signaling and repair factors ^1^. Thus, we expected our hitlist to include genes that have a described role in the RS-response and the DDR^36–38^. Indeed, we identified multiple genes previously shown to be involved in the RS-response/DDR, including multiple *FANC* genes ^39,40^ *LIG3*^44–46^, *UBR5*^47–49^, *and BRIP1*^42,43,50,51^ and (Fig. 2B-D). This indicates that our screen was successful in identifying vulnerabilities of cells experiencing RS. In addition to the identification of anticipated factors, we also identified several hits with no previous link to the RS-response or DDR, including a set of genes related to mitochondrial function (Fig. 2C, D).

### Mitochondrial genes and the RS-response

Interestingly, amongst the hits there was a strong enrichment of genes related to mitochondrial processes (Fig. 2C,D). It is unlikely that this response is due to the induction of replication stress in mitochondria specifically as Aphidicolin only interferes with replication of the nuclear DNA^51^. To confirm the synthetic lethal interaction between mitochondrial genes and RS in our screens, we selected POLG2, one of the most prominent mitochondrial-associated hits, for further validation (Fig. 2B). siRNA-mediated depletion of POLG2 did not result in increased sensitivity to Aph or HU in growth assays, when compared to WT HAP1 cells (Fig. S2A/B). Although knockdown was relatively prominent (>90%) (Fig. S2C), it is possible that residual POLG2 was sufficient to sustain mitochondrial function. To eliminate this possibility, we extensively attempted to generate KO clones for multiple mitochondria-related hits using CRIPSR/Cas9 but this was unsuccessful. As an alternative approach to interfere with mitochondrial function, we inhibited oxidative phosphorylation, a key process of mitochondria, using oligomycin. We selected a dose of 2.5 μM oligomycin that only mildly affected growth (data not shown). However, when combining oligomycin with Aph or HU treatment, no enhanced sensitivity could be observed (Fig. S2D).

Even though we were unable to validate the synthetic lethal interaction between loss of mitochondrial genes and RS as observed in our screens, the prominent enrichment of mitochondrial hits remains interesting. Further validation of these hits is required to explore a potential link between mitochondrial function and RS.

### Validation of novel genes in the RS response

Next, we set out to confirm the hits with no previous role in the RS response. For this, we systematically generated knock-outs of the top hits from our screen using CRISPR-Cas9 (see **methods** for details). We excluded all mitochondrial related hits and selected a subset of hits with no or limited described roles in the RS response. We successfully generated one or more knock-out clones for *RNASEH1, HNRNPA2B1, VEZT, GIGYF2, SUMO2, and CDC42*.

Next, we subjected the different KO cell-lines to growth assays in the presence of increasing doses of Aph or HU (Fig. 3). We found that all knock-out cell lines showed increased sensitivity to Aph and HU treatment, as evidenced by the decrease in IC50 compared to WT cells (Fig. 3). This shows that our screen obtained some novel candidates with potential roles in the RS response.

**Figure 3:**
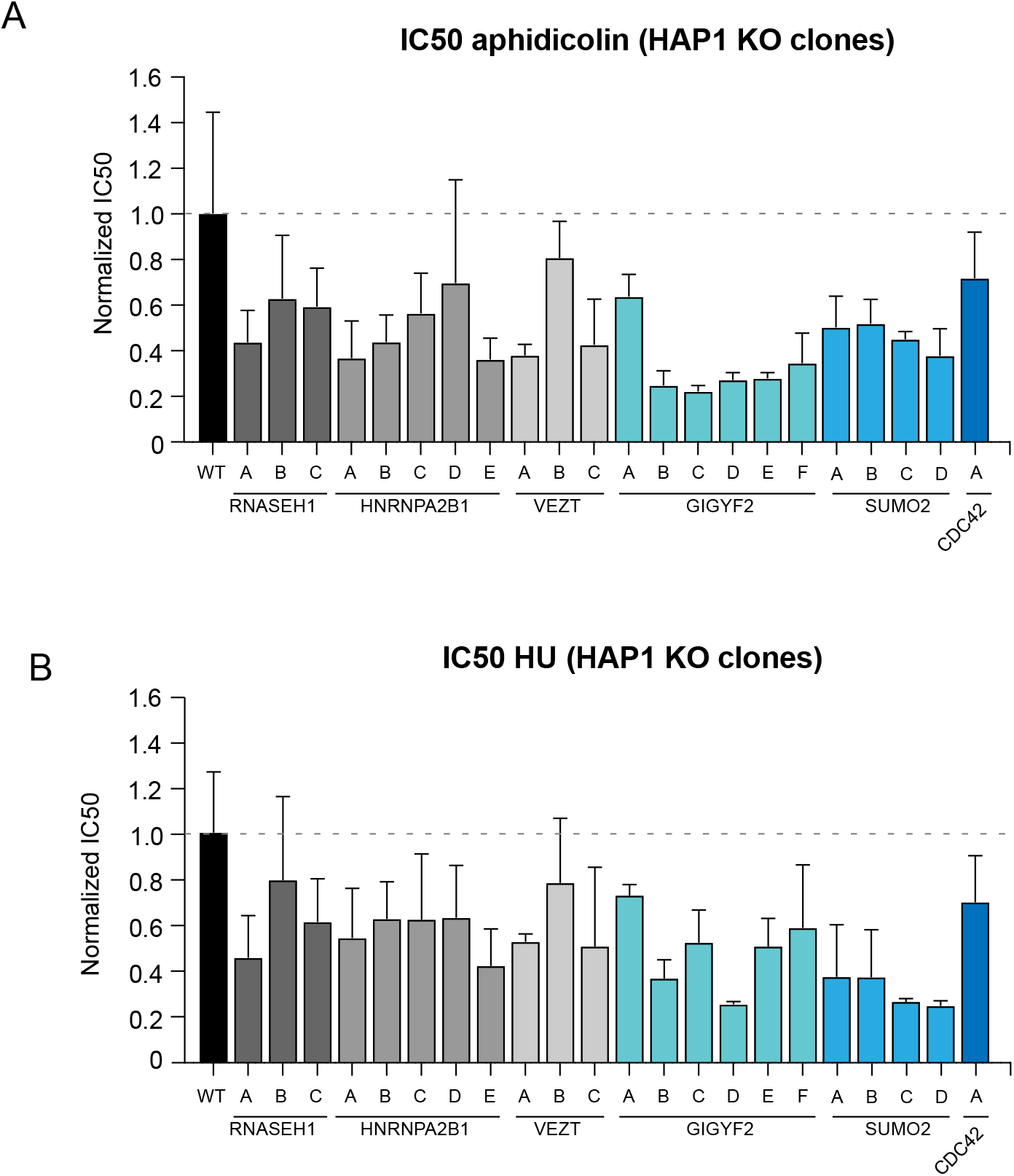
Sensitivity to replication stress could be confirmed for multiple hits by CRISPR-mediated gene knock-outs. **(A)** Average IC50 of growth assays for WT HAP cells and different knock-out cell-lines subjected to Aph. At least 2 replicates per knockout were performed. **(B)** Average IC50 of growth assays for WT HAP cells and different knock-out cell-lines subjected to HU. At least 2 replicates per knockout were performed.

### GIGYF2 might be involved in HR-mediated DNA-damage repair

Out of our validated hits, GIGYF2 was one of the most prominent hits and sensitivity to RS could be confirmed in 6 independent KO clones, for both HU and Aphidicolin (Fig 3). Therefore, we selected GIGYF2 for further validation. GIGYF2 is a key component of the 4EHP-GYF2 multiprotein complex involved in repression of the initiation of translation^52^, and no role in the RS or DDR response has been described before.

We selected clone B for further analysis and confirmed full loss of the protein by western blot analysis (Fig 4A). Growth assays for this specific clone confirmed the enhanced sensitivity to both Aphidicolin and HU (Fig. 4B). IC50 values were calculated using nonlinear regression analysis and averaged from minimally 3 independent experiments (Fig. 4C). Interestingly, GIGYF2 was recently also identified in a haploid genetic screen in cells subjected to ionizing radiation (IR)^53^ (Fig. 4D). Indeed, we confirmed that ΔGIGYF2 cells displayed increased sensitivity to IR compared to WT HAP1 cells (Fig. 4E). To test whether GIGYF2 has a role in DNA damage repair, we studied the double stranded break (DSB) repair kinetics of WT HAP1 and ΔGIGYF2 cells through measuring the amount of yH2AX foci at several time-points after irradiation (IR) (Fig. 4F). Besides a baseline increase in the level of yH2AX foci in ΔGIGYF2 cells compared to WT HAP1 cells (Fig. 4F, t=0), we also observed a prominent delay in repair (Fig. 4F, t=3h post IR). This implies that ΔGIGYF2 cells have a defect in DNA damage repair which might explain their enhanced sensitivity to RS.

**Figure 4:**
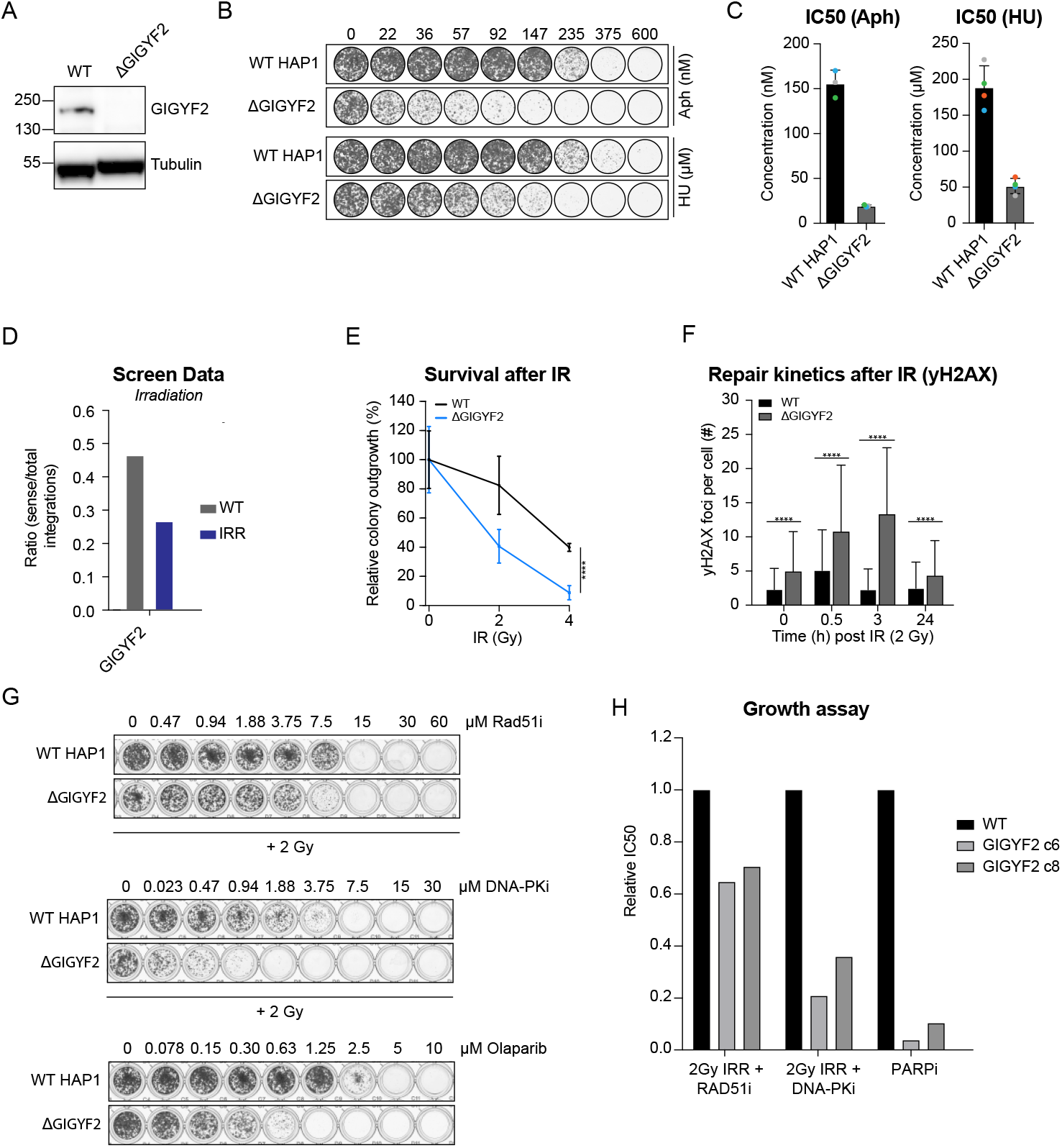
Loss of GIGYF2 sensitizes cells to IR and leads to impaired DNA damage repair. **(A)** Western blot confirming knock-out of GIGYF2. Tubulin serves as loading control. **(B)** Representative growth assays of WT HAP1 and ΔGIGYF2 HAP1 cells, subjected to increasing doses of Aph or HU for 5 days. **(C)** Average IC50 of growth assays, where WT HAP1 and ΔGIGYF2 HAP1 cells were subjected to Aph or HU (n=3). **(D)** Bar graph of the average ratio of sense/total integrations of GIGYF2 in WT, and (every other day) 1 Gy irradiated HAP1 cells. The data are extracted from 4 individual WT insertional mutagenesis screens, and 1 individual IR insertional mutagenesis screen. **(E)** Relative outgrowth capacity of WT and ΔGIGYF2 HAP1 cells after increasing doses of IR, average of 4 independent experiments. Mean + SD, ** p<0.01, **** p<0.0001 (unpaired students t-test). **(F)** Number of yH2AX foci in asynchronous WT and ΔGIGYF2 HAP1 cells at indicated time points after IR, average of minimally 200 cells per condition, of 1 independent experiment. Mean + SD, **** p<0.0001 (unpaired students t-test). **(G)** Growth assays of WT HAP1 and ΔGIGYF2 HAP1 cells, subjected to increasing doses of RAD51 inhibitor for 5 days (cells plated in inhibitor, before being subjected to 2 Gy of IR on day 0), increasing doses of DNA-PK inhibitor for 5 days (cells plated in inhibitor, before being subjected to 2 Gy of IR on day 0), or increasing doses of olaparib for 5 days. IC50 values displayed on the right corresponding to the cell-line. **(H)** Bar graph displaying the IC50 values from the growth assays (4F).

To test in which repair pathway GIGYF2 plays a role, we performed growth assays in the presence of increasing doses of drugs targeting specific DNA-repair pathways. By inhibiting one repair pathway, cells become more dependent on alternative pathways for DSB repair. For example, when inhibiting non-homologous end joining (NHEJ), cells rely more on homologous recombination (HR) to resolve a DSB^54^. We used DNA-Pk inhibition (DNA-Pki) to inhibit NHEJ^55^, Rad51 inhibition (Rad51i) to inhibit HR^56^, or PARP trapping (olaparib), which is selectively more toxic to HR-deficient cells^57–59^. DNA-PKi and Rad51i were combined with 2 Gy IR to generate DSBs and activate the DDR. Strikingly, we found that ΔGIGYF2 cells showed a mild increase in sensitivity to Rad51i + 2 Gy (Fig. 4G/H). More strikingly, ΔGIGYF2 cells displayed a dramatic increased sensitivity to DNA-Pki + 2 Gy (Fig. 4G/H) and to olaparib (Fig. 4G/H), behavior that is consistent with a defect in HR. Altogether, our results suggest that GIGYF2 is important under conditions of RS, likely through a function in HR-mediated repair. HR is important for the repair of stalled or broken forks and protects from genomic instability caused by RS. The exact role of GIGYF2 in this pathway requires further investigation.

### SUMO2, but not SUMO3, might be important for HR-mediated DNA damage repair

We next set out to validate and investigate the specific dependency on SUMO2 of cells experiencing RS. Small Ubiquitin-like Modifier (SUMO) proteins are small proteins with a high similarity to ubiquitin. The modification of proteins by SUMO (referred to as SUMOylation) can affect the targets stability, localization or functionality. Human cells express 5 SUMO proteins but there are 3 main players: SUMO1, SUMO2 and SUMO3^60^. SUMO1 is responsible for mono-SUMOylation and shares about 50% sequence similarity with SUMO2 and SUMO3. In contrast, SUMO2 and SUMO3 share 97% sequence identity with each other and are known to generate longer sumo chains. Since SUMO2 and SUMO3 are structurally extremely similar, they are generally considered functionally redundant. Hence, they are often referred to as SUMO2/3 and not separately. Although SUMO2 and SUMO3 are both prominently expressed in HAP1 cells^61^, it was surprising that we found a very specific role for SUMO2 in the RS screens, while SUMO3 seemed to completely dispensable (Fig. 5A). This suggests that SUMO2 and SUMO3 are not completely redundant, at least not in the context of RS, which prompted us to investigate this in more detail.

**Figure 5:**
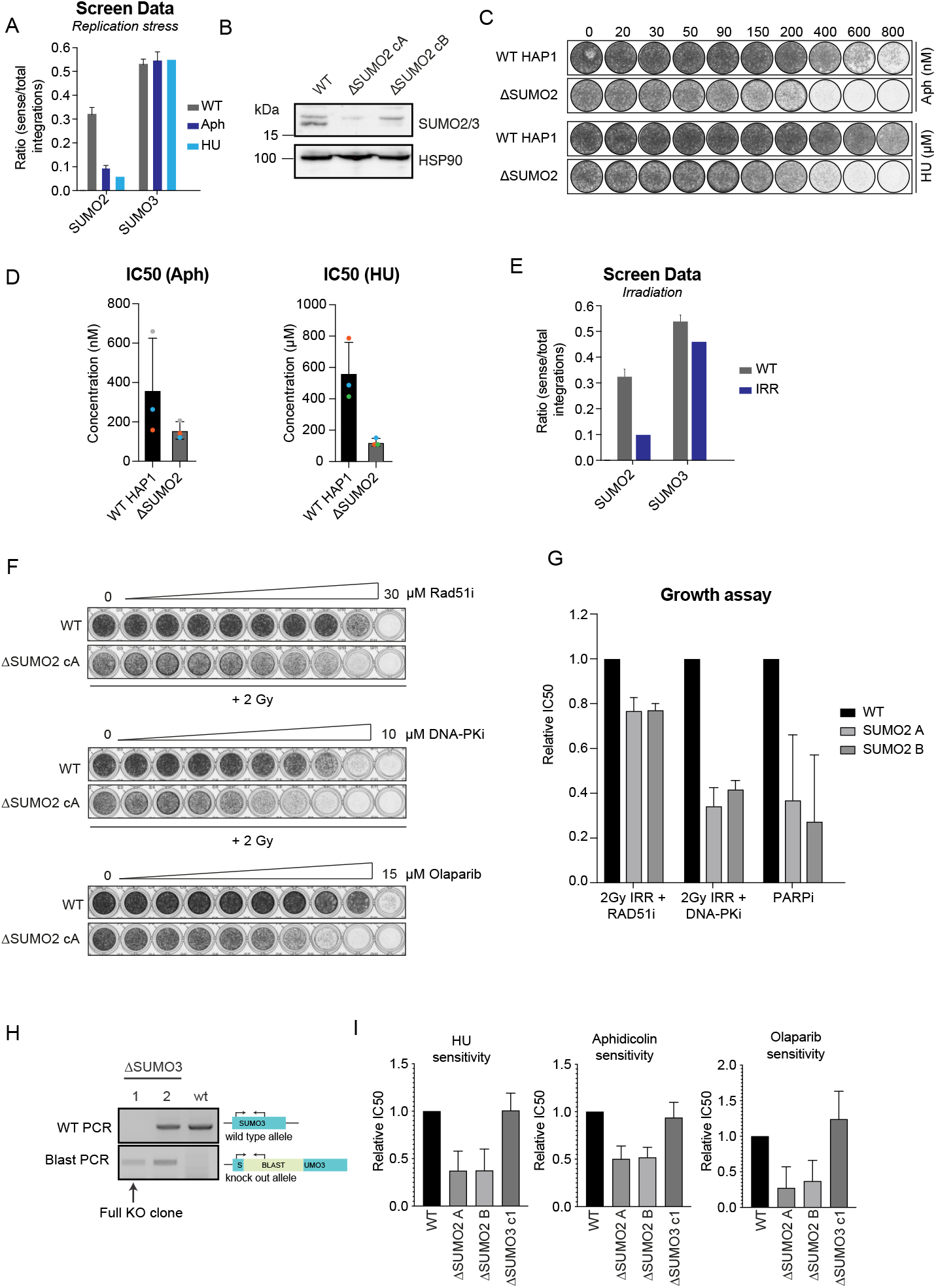
Loss of SUMO2, but not SUMO3, sensitizes cells to DNA damage. **(A)** Bar graph of the average ratio of sense/total integrations of SUMO2 and SUMO3 in WT, and Aph/HU treated HAP1 cells. The data are extracted from 4 individual WT insertional mutagenesis screens, 2 individual Aph insertional mutagenesis screens, or 1 individual HU insertional mutagenesis screen. **(B)** Western blot confirming knock-out of SUMO2. HSP90 serves as loading control. **(C)** Representative growth assays of WT HAP1 and ΔSUMO2 HAP1 cells, subjected to increasing doses of Aph or HU for 5 days. **(D)** Average IC50 of growth assays, where WT HAP1 and ΔSUMO2 HAP1 cells were subjected to Aph or HU (n=3). **(E)** Bar graph of the average ratio of sense/total integrations of SUMO2 and SUMO3 in WT, and (every other day) 1 Gy irradiated HAP1 cells. The data are extracted from 4 individual WT insertional mutagenesis screens, and 1 individual IR insertional mutagenesis screen. **(F)** Growth assays of WT HAP1 and ΔSUMO2 HAP1 cells, subjected to increasing doses of RAD51 inhibitor for 5 days (and subjected to 2 Gy of IR on day 0), increasing doses of DNA-PK inhibitor for 5 days (and subjected to 2 Gy of IR on day 0), or increasing doses of olaparib for 5 days. **(G)** Bar graph displaying the IC50 values from the growth assays (5F). **(H)** PCR to identify integration of a blasticidin-resistance cassette in the SUMO3 gene. Clone 1 and 2 show successful integration but only clone 1 shows the disruption of the WT allele. Therefore, clone 1 was selected for further validation experiments. confirm SUMO3 knock-outs. **(I)** Bar graphs displaying the IC50 of WT HAP1, ΔSUMO2 HAP1, and ΔSUMO3 HAP1 cells subjected to either Aph, HU or olaparib.

We first set out to validate and investigate the role of SUMO2 in RS. Indeed, HAP1 clones that were deleted for SUMO2 (Fig. 5B) displayed enhanced sensitivity to both HU and Aphidicolin (Fig. 5 C/D). To test if SUMO2 has a role in double stranded break repair, we consulted the data from the synthetic lethality screen for ionizing irradiation^53^. Strikingly, the screen results suggest that SUMO2 loss also sensitizes HAP1 cells to double stranded breaks induced by irradiation (Fig. 5E). Also in this context, SUMO3 was dispensable. To test if SUMO2 has a role in a specific repair pathway, we again treated HAP1 cells with specific repair inhibitors for NHEJ (DNA-Pki), HR (RAD51i) or with the PARP trapping agent Olaparib. DNA-PKi and Rad51i were combined with 2 Gy IR to generate DSBs and activate the DDR. We found that ΔSUMO2 cells showed only a mild increase in sensitivity to Rad51i (Fig. 5 F/G). More strikingly, ΔSUMO2 cells were considerably more sensitive to DNA-PKi and to Olaparib (Fig. 5 F/G). This behavior is consistent with a defect in HR. To make sure that these functions are specific to SUMO2, we also generated a SUMO3 KO clone. Since no SUMO3 specific antibody exists, we could not confirm loss of the protein on western blot. Therefore, we tested for the successful integration of a blasticidin-resistance cassette by PCR on genomic DNA (Fig. 5H). Clone 1 shows disruption of the WT-allele concomitant with successful integration of the resistance cassette. Consistent with the screen data, SUMO3 KO clone 1 did not sensitize to HU and Aphidicolin, and also not to Olaparib (Fig. 5I). These results further implicate a role for SUMO2, and not SUMO3, in HR.

## Discussion

Replication stress (RS) is a severe threat to genomic stability, and highly evident in (pre-) cancerous lesions^1,11^. Cells have evolved mechanisms to deal with RS and resolve issues that originate from prolonged fork stalling, such as checkpoint activation and activation of DNA repair pathways^1,5,6^. Since RS is a cancer cell-specific feature, targeting the RS-response has been of therapeutic interest (reviewed by Ubhi et al.^16^ and Zhang et al.^7^). The rationale being that the inhibition of key players of the RS-response in cancer cells with high baseline levels of RS will increase genomic instability to levels that induce cancer-specific cell killing^37^.

In this work, we set out to discover genes that are specifically essential for cells enduring RS, using a well-established haploid genetic screening approach^35^. We identified genes whose loss compromises proliferation of cells treated with RS-inducing drugs, while minimally affecting untreated cells. Besides the identification of genes with a well described role in the RS-response/DDR, we also discovered a set of genes for which no function in replication/DDR had been described thus far. First, we discovered a large set of mitochondrial-related genes. Mitochondria are key regulators of cellular metabolism, their most prominent role being the production of ATP through oxidative phosphorylation (OXPHOS)^62^. Previously, it has been proposed that energy adaptation through mitochondria is important under conditions of cellular stress^63,64^. We aimed to validate and test this hypothesis in light of RS, but we were unable to generate knockouts for the mitochondrial genes. Therefore, we propose to use acute depletion, for example by using an auxin-inducible degron or by exploiting a CRISPRi system to interfere with these genes. These approached could help to validate our RS screen before investigating if the mitochondria indeed create an environment or energy state that helps cells to cope with RS.

Importantly, we could generate knockout cells and validate that *GIGYF2, SUMO2, HNRNPA2B1, RNASEH1, VEZT*, and *CDC42* are specifically required in cells suffering from RS. We did not resolve the full mechanism by which these proteins function in the RS-response but we will speculate on their potential roles below. GIGYF2 is involved in repressing the initiation of translation^52^, and has been shown to prevent formation of toxic protein intermediates from ribosomes trapped with faulty mRNAs^65^. We found that ΔGIGYF2 cells are more sensitive to Aph, HU and IR, display an increase in DNA damage at baseline, and decreased repair kinetics compared to WT HAP1 cells (Fig. 4). This suggests that GIGYF2 aids in reducing genomic instability, likely through a role in DNA damage repair. Importantly, ΔGIGYF2 cells showed increased sensitivity to PARP inhibition and to inhibition of NHEJ (Fig. 4C, D). This behavior resembles the synthetic lethal interaction of PARP inhibition with HR-deficiency, such as found in BRCA1/2 mutated tumors^57–59^. Interestingly, a genome-scale CRISPRi-Cas9 screen has identified GIGYF2 to be selectively essential in RPE-1 cells harboring a CCNE1 overexpression construct, a common trait of cancer cells that results in enhanced levels of RS and genomic instability ^66^. Furthermore, genomic analysis of ovarian cancers found CCNE1 amplification to be mutually exclusive with HR-deficiency^67,68^. These findings further highlight the link between the RS-response and HR-deficiency and further suggest a potential role for GIGYF2 in HR. Further work is needed to identify the exact role of GIGYF2 in HR. For example, GIGYF2 could have a direct role during the repair of DNA damage lesions. Alternatively, due to the described role of GIGYF2 in translational repression^52^, it is also possible that its role is more indirect, for example by causing DDR-protein imbalances.

SUMO proteins aid in a range of biological processes, including DSB repair, through post-translational modification of a plethora of proteins^69–71^. The number of proteins regulated by SUMOylation in the context of DNA damage is likely incomplete as new targets of SUMO are still emerging. For example, a recent publication described the SUMOylation of HNRNPA2B1, another hit of this screen (Fig 2 and 3), to be critical for a functional HR pathway^72^. Our data also hint towards a role for SUMO2 in HR. Such a role is not unexpected, as multiple proteins in the HR pathway are described to be regulated by SUMOylation. For example, resection by CtIP is regulated by its SUMOylation^73^, Mdc1 has been shown to be SUMOylated by PIAS4 to promote its turnover^74^ and modification of PML by SUMOylation facilitates the formation of PML nuclear bodies^75^. It would be interesting to further investigate the role of SUMO2 in HR, for example by comparing SUMOylation of HR proteins in WT HAP1 cells and ΔSUMO2 cells.

We specifically picked up SUMO2 to be essential during RS, and not SUMO3. This is surprising as SUMO2 and SUMO3 are thought to be functionally redundant^71^. It is possible that this specificity can be attributed to unique features of SUMO2. For example, it has been described that SUMO2 has a different localization pattern as compared to SUMO3^76^. However, it is also possible that the different dependencies can be attributed to their relative expression levels. During mouse embryonic development, SUMO2 is essential while SUMO3 is dispensable^77^. This is explained by the relative high expression of SUMO2 during development compared to SUMO3. It is therefore not unlikely that SUMO2 is simply more essential in the context of our screens because the relative expression levels of SUMO2 and SUMO3 is also different in the context of HAP1 cells (Fig 5B). If expression level differences indeed explain the different dependency on SUMO2 versus SUMO3, it would be sufficient to replenish the low levels of SUMO in the SUMO2 KO cells with excess SUMO3. It would be also interesting to investigate if the specific dependency on SUMO2 also exists in other contexts and processes where sumoylation plays an important role, such as in metabolism, nuclear transport and RNA processing^60^.

Altogether, we believe our work could be a valuable resource of RS-specific vulnerabilities. We confirm that cells under RS depend on a proper DDR, evident by the essentiality of many DDR-related genes in our RS-screen. This result compliments strategies exploited in the clinic, that aim to eradicate cancer cells by exacerbating their levels of genomic instability through for example ATR inhibition^37,38^. It is important to note that the level and type of RS in cancer cells may differ from the level of RS obtained in our screens by treating HAP1 cells with low levels of specific RS-inducing drugs. HAP1 cells are a derivative from the chronic myeloid leukemia cell line KBM7^35^, and display basal levels of RS. We therefore cannot exclude the possibility that the identified hits only become essential when further enhancing RS in cancer cells that already display baseline RS or that the interaction is specific to the RS inducing drugs. It would be interesting to systematically test the essentiality of our screen hits in an array of cancer cell lines with varying levels of RS as well as some untransformed cell lines. Moreover, it would be interesting to test the sensitivity of cell lines that are deficient for the different hits against a large panel of DNA damage inducing drugs and repair inhibitors. Such systematic approaches could uncover the precise cellular contexts in which these hits become essential for viability and give important insights into the pathways these hits act to aid in the tolerance towards RS.

## Methods

### Cell culture

Human derived near-haploid HAP1 cells were cultured in IMDM (GIBCO) supplemented with 10% FCS, 1% GlutaMAX supplement (GIBCO), 100 U/ml penicillin, and 100 μg/ml streptomycin. siRNA transfections were performed using RNAiMax (Invitrogen) according to the manufacture’s guidelines. The following siRNAs were used in this study: siNon-Targetable (Dharmacon), siPOLG2 (Horizon, ON-TARGETplus, SMARTPool), siMRPL23 (Horizon, ON-TARGETplus, SMARTPool). All drugs (Aphidicolin, HU, olaparib, RAD51i (B02), DNA-PKi (NU7441), and oligomycin A) were dissolved in DMSO and used at indicated concentrations. Cells were γ-irradiated using a Gammacell Extractor (Best Theratronics) with a 137Cs-source.

### Growth assays

HAP1 cells were plated in 96-well plates, at a density of 1500 cells/well and treated as indicated for 5 days. After 5 days, cells were fixed using 100% methanol, and stained for 2h at room temperature using crystal violet. Subsequently, crystal violet was dissolved in 10% acetic acid, and intensity was measured using a BioTek Epoch Spectrophotometer at 595 nm. These measurements were used for IC50 calculations in PRISM, using nonlinear fit, sigmoidal, 4PL, X is log (concentration).

### Immunofluorescence

Cells were grown on 9mm glass coverslips and fixed for 10 minutes at room temperature in 4% formaldehyde with 0.2% Triton X-100. The following antibodies were used: human anti-Crest (Cortex Biochem, cs1058), rabbit anti-pH3Ser10 (Campro, #07-081), mouse anti-ERCC6L (PICH) (Abnova, 000548421-b01p). All primary antibodies were incubated over night at 4°C. Secondary antibodies (Molecular probes, Invitrogen) and DAPI were incubated for 2 hours at room temperature. Coverslips were mounted using ProLong Gold (Invitrogen). Images were acquired using a Deltavision deconvolution microscope (Applied Precision) with a 60x 1.40 NA oil objective. Softworx (Applied Precision), ImageJ, Adobe Photoshop and Illustrator CS6 were used to process acquired images.

### Haploid insertional mutagenesis screens

Genes essential for survival of HAP1 cells treated with either Aph or HU were identified using a haploid insertional mutagenesis screen as described previously^35^. Mutagenized HAP1 cells were obtained from the Brummelkamp laboratory. In brief, mutagenesis of HAP1 cells was obtained as follows: gene trap retrovirus was produced in HEK293T cells. Retrovirus was harvested twice daily for a minimum of three days, and pelleted by centrifugation (2 hours, 21,000 rpm, 4°C using a SW28 rotor). Approximately 40 million HAP1 cells were mutagenized by transduction of the concentrated gene trap virus in the presence of 8 μg/ml protamine sulfate in a T175 flask for at least two consecutive days. The mutagenized cells were frozen in IMDM medium containing 10% DMSO and 10% FCS. After thawing, mutagenized HAP1 cells were passaged for 10 days in the presence of either 27.5 nM aphidicolin or 100 μM HU. After passaging, cells were collected by trypsin-EDTA followed by pelleting. Cells were fixed using fix buffer I (BD biosciences). To minimize confounding from diploid cells potentially harboring heterozygous mutations, fixed cells were stained with DAPI to allow sorting on G1 haploid DNA content using an Astrios Moflo. 30 million sorted cells were lysed overnight at 56°C to allow for de-crosslinking followed by genomic DNA isolation using a DNA mini kit (QIAGEN).

### Insertion site mapping

The gene trap insertion sites were amplified by LAM-PCR, followed by capture, ssDNA linker ligation, and exponential amplification using primers containing illumina adapters prior to sequencing, as described previously ^35^. Mapping and analysis of insertions sites is described previously^78^. Briefly, following sequencing on a HiSeq 2000 or HiSeq 2500 (Illumina), insertion sites were mapped to the human genome (h19) allowing one mismatch, and intersected with RefSeq coordinates to assign insertions sites to genes. Gene regions overlapping on opposite strands were not considered for analysis, while for genes overlapping on the same strand gene names were concatenated. For each replicate and for both drug treatments (Aph or HU) gene essentiality was determined by binomial test. Synthetic lethality was assessed by comparing the distribution of sense and antisense orientation integrations for each gene in the drug treated replicates with 4 wild-type control datasets previously published^79^ (NCBI SRA accession number SRP058962) using Fisher’s exact tests. A gene was considered a screen specific hit when it passed all Fisher’s tests with a P value cutoff of 0.05 and an effect size of at least 0.12 (subtraction ratio WT sense ratio – replication stress condition sense ratio).

The final hit list is composed of genes that met the selection criteria in the HU screen and at least in one of the Aphidicolin replicates, to find general replication stress sensitizers, and exclude treatment specific hits.

### CRISPR-mediated generation of knockout cell-lines

Knockout cells were generated using CRISPR/Cas9 mediated genome editing. Guide sequences were designed using CRISPOR design. Guides to generate knock-out cell-lines were targeted against exon 1, 2 or 3 of the gene of interest, and subsequently cloned into the pX330 vector (Addgene plasmid #42230). pX330 and a donor vector containing a blasticidine resistance-cassette^80^ were co-transfected in HAP1 cells and selected with 10 μg/ml blasticidin. Individual clones were selected and knock-outs were confirmed using PCR to confirm integration of the blasticidin cassette at the correct locus, and by western blot analysis where indicated. The following guides were used:

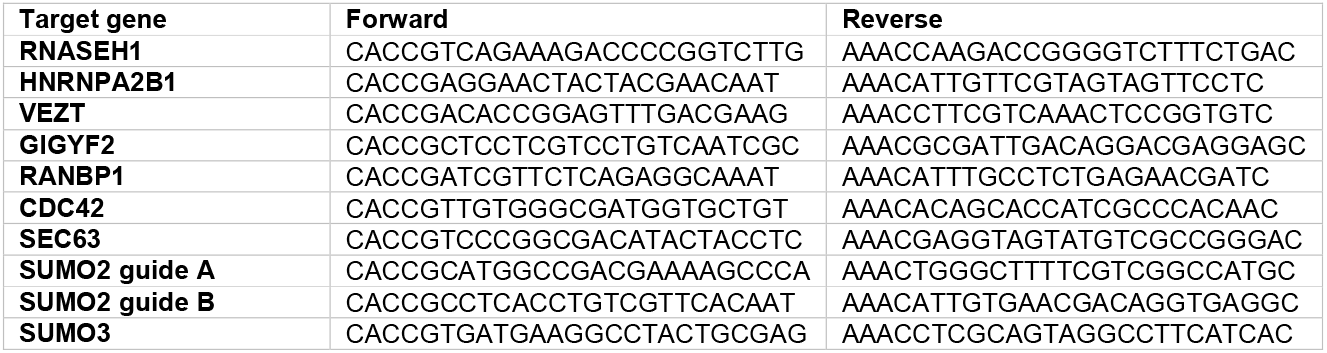

### RNA isolation and qRT-PCR

Cells were harvested by trypsinization 4 days post siRNA-transfection. RNA was isolated using the RNeasy kit (Qiagen) and quantified by NanoDrop (Thermo scientific). cDNA was synthesized using Bioscript Reverse Transcriptase (Bioline), random primers (Promega), dNTPs, and 800 ng of total RNA, according to manufacturer’s protocol. Primers were designed with a Tm close to 60 °C to generate 90-120 bp amplicons, spanning introns. Subsequently, 10 ng of cDNA was amplified for 40 cycles on a Roche Lightcycler 480, using SYBR Green PCR Master Mix (applied Biosystems). Target cDNA levels were analyzed by comparative cycle (Ct) method and values were normalized to GAPDH expression levels. The following primers were used:

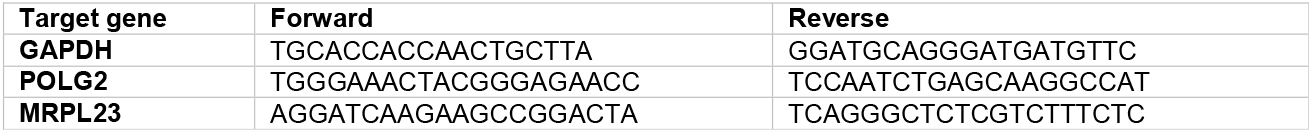

### Western-blot

Cells were lysed using Laemmli buffer (120 mM Tris pH 6.8, 4% SDS, 20% glycerol), and treated with drugs as indicated. Equal amounts of protein were separated on a polyacrylamide gel followed by transfer to a nitrocellulose membrane. Membranes were blocked in a 5% milk/TBS solution. Antibodies were incubated in 2,5% milk in TBS containing 0,1% Tween. The following antibodies were used in this study: mouse anti-alpha-tubulin (Sigma, t5168), rabbit-anti-HSP90 (Santa Cruz, sc7947), mouse anti-GIGYF2 (Santa Cruz, sc393918), rabbit anti-SUMO2/3 (Santa Cruz, SC-32873). HRP-coupled secondary antibodies (DAKO) were incubated for 2h at room temperature in a 1:2500 dilution. The immune-positive bands were visualized using Immobilon Western HRP Substrate (Millipore) and a ChemiDoc MP System (Biorad).

### Quantification and Statistical Analysis

Significant differences between treatment conditions were calculated using a Student’s t-test. In all figures: ∗, p value < 0.05; ∗∗, p value < 0.01; ∗∗∗, p value < 0.001; ∗∗∗∗, p value < 0.0001. Gene set enrichment analysis was performed using ClusterProfiler (version 4.6.2) in R. Correlation of screen data was calculated using Pearson’s correlation coefficient.

## Acknowledgments

We would like to thank the R.H. Medema, T. Brummelkamp, B. Rowland and J. Jacobs laboratories for helpful discussions. This study was supported by funds from the Dutch Cancer Society (KWF-NKI-2015-7832), granted to R.H.M. and J.A.R.

