## Supplemental Figures 1-2 for "Identification of critical factors of the replication stress response in human cells"

### Fig S1: Correlations between screens

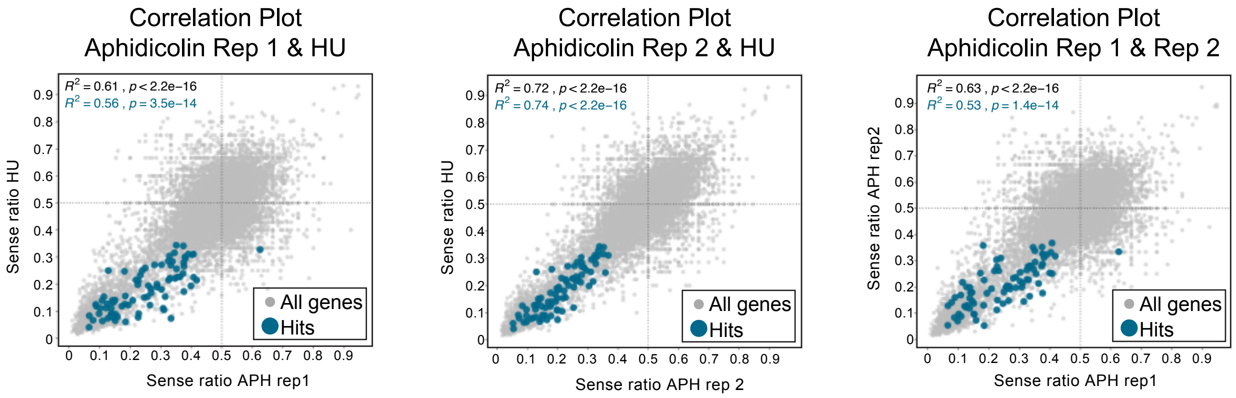

Fig S2: Depletion of POLG2 does not sensitize HAP1 cells to replication stress.

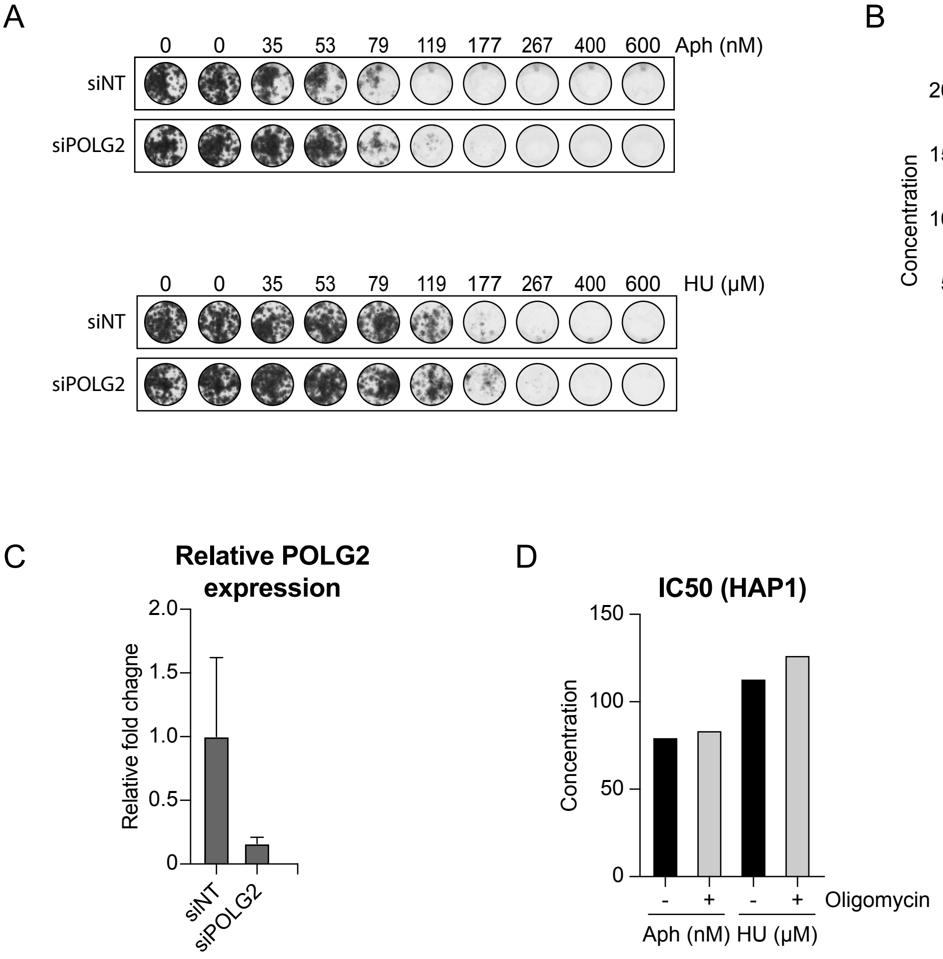
